# Müller glia-vasculature interactions in the developing retina

**DOI:** 10.64898/2026.03.23.713726

**Authors:** Samira Monshietehadi, Angel J. Garcia, Benjamin Smith, Matthew Tarchick, Aileen Conley, Kiara Abhayaratne, Marla B. Feller

## Abstract

Coordinated signaling among neurons, glia, and the vasculature is essential for the formation of a functional nervous system, yet how these relationships emerge during development remains unclear. Here, we investigated the developmental interplay between neural activity, Müller glia, and the retina vasculature in mice. Using quantitative confocal imaging from postnatal day 5 to eye-opening, we mapped the emergence of the superficial, intermediate, and deep vasculature layers and found that they emerged normally in mice lacking the β2-containing nicotinic acetylcholine receptors, despite a dramatic reduction in cholinergic signaling. Tip cell density and overall vessel growth were unchanged, indicating cholinergic wave activity is not required for the emergence of retinal vasculature.

We next defined the developmental timeline of Müller glia–vascular interactions. Sparse labeling and immunohistochemistry revealed that Müller glial lateral processes closely associate with endothelial tip cells during intermediate- and deep-layer angiogenesis and establish Aquaporin-4–enriched endfeet at vascular contact sites from the earliest stages of growth, even when vessel trajectories are perturbed.

Finally, two-photon calcium imaging combined with simultaneous electrophysiology demonstrated that Müller glial endfeet exhibit robust, compartmentalized calcium transients during development. Although a subset of events was temporally correlated with retinal waves, enhancing neurotransmitter spillover selectively increased wave-associated activity in glial stalks but not endfeet. These findings indicate that calcium signaling at the glial-vascular interface is largely independent of spontaneous neuronal activity. Together, our results support a model in which Müller glia engage growing vessels through an activity-independent, parallel developmental program that may provide instructive cues for retinal angiogenesis.

## Introduction

During development, signaling between neurons, glia, and blood vessels is crucial for constructing a healthy nervous system (Wälchli et al. 2023; Biswas et al. 2020). This coordination is especially critical in the retina, where disruptions in vasculature growth underlie vision loss in conditions such as retinopathy of prematurity (Zhang et al. 2023; Selvam et al. 2018). Although molecular pathways that guide angiogenesis have been well characterized (Ferrara 2004; Fruttiger 2007; Hellström et al. 2007), growing evidence suggests a role for neural activity and glial signaling in the development of CNS (Lacoste et al. 2014; Whiteus et al. 2014; Iadecola 2017; Rattner et al. 2019; Matsuoka et al. 2016).

The vasculature in the retina is comprised of two distinct vascular beds—the choroidal vasculature, which supports the metabolically active pigment epithelium and photoreceptor outer segments, and the retinal vasculature, which oxygenates the neural retina in mammals with thick retinas. The retina vasculature is organized into three planar layers (Figure 1A): (1) the superficial plexus of arteries, veins, and an extensive capillary bed that resides on the surface of the retina in the retinal nerve fiber layer; (2) the deep capillary bed at the outer nuclear layer/outer plexiform layer border (ONL/OPL); and (3) the intermediate capillary bed at the inner nuclear layer/inner plexiform layer (INL/IPL). All three layers are connected by perpendicular diving blood vessels.

**Figure 1:**
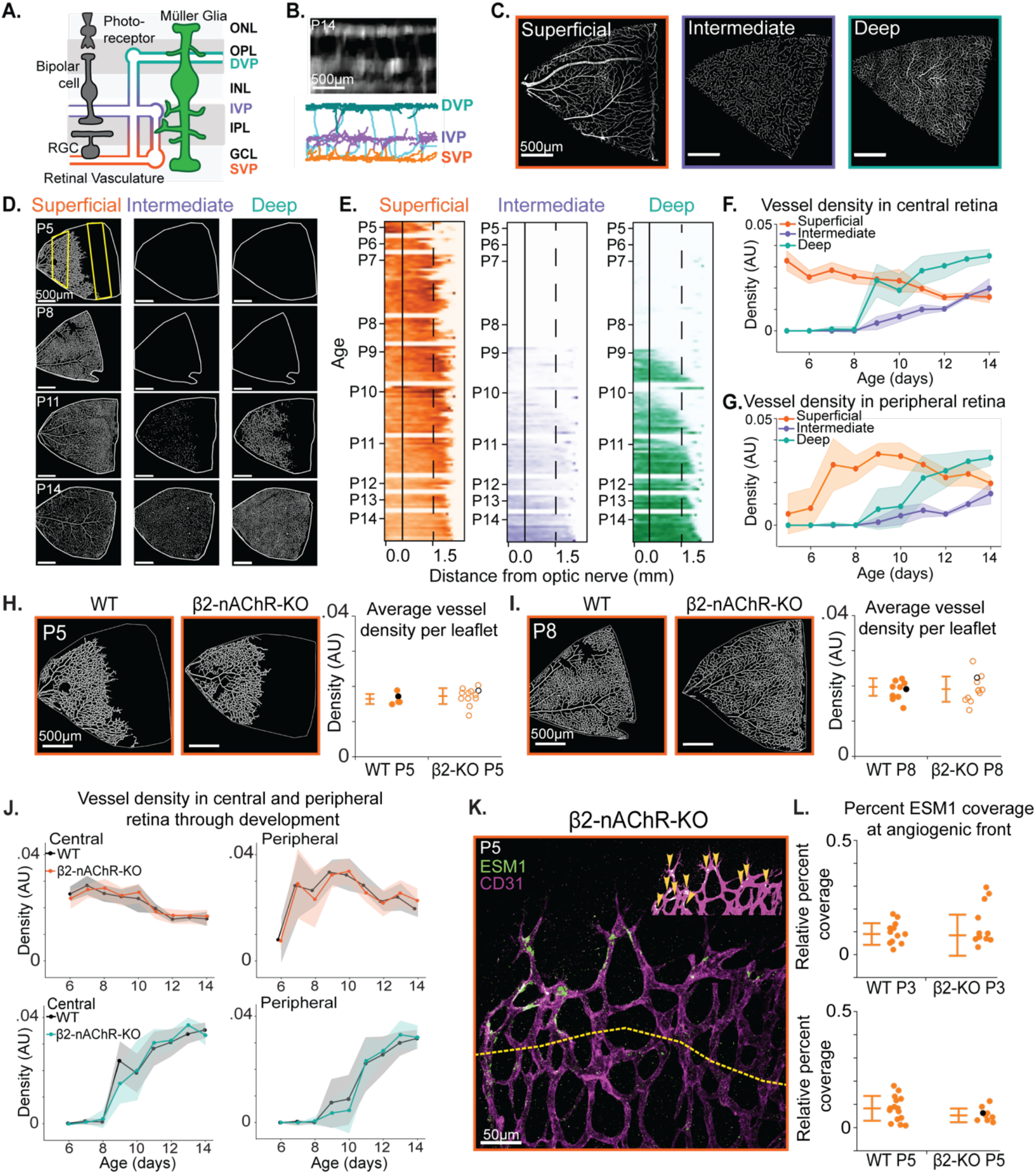
Retina vasculature grows uniformly throughout development and grows independently of cholinergic retinal waves. A. A schematized cross-section view of the retina: neurons (black), glial cells (green) and the superficial (SVP, orange), intermediate (IVP, purple), deep layer (DVP, teal). B. Top: A cross-section view of a P14 retina vasculature labeled with CD31 to view diving vessels. Scale = 500μm. Bottom: Skeletonized vasculature path. C. Example images of a single leaflet of developed superficial, intermediate, and deep retinal vasculature layers. Vasculature labelled using antibody to CD31 and imaged using a tile-scanning confocal microscope. Scale = 500μm. D. Example images of superficial, intermediate and deep vasculature layers at different points during development. Vasculature labelled using antibody to CD31 and imaged using a tile-scanning confocal microscope. Scale = 500μm. E. Quantification of D. Density as a function of distance from the optic nerve was computed using a sliding 100μm wide window (yellow box in D) from the optic nerve to the distal end of the retina leaflet. Each line corresponds to one leaflet. Deeper color corresponds to denser vessels in that region. F. Plot of center region 0.4mm from the optic nerve (solid line in E) showing vessel density as a function of age for the superficial, intermediate, and deep layers. G. Plot of peripheral region 1.6mm from the optic nerve (dashed line in E) showing vessel density as a function of age for the superficial, intermediate, and deep layers. H. Left: Representative superficial layer images of a P5 WT and β2 KO retina flat mounts stained for endothelial cell marker CD31. Scale bar = 500μm. Right: Average vessel density across each leaflet at P5. Each point represents 1 leaflet. Points are acquired by averaging all 100μm windows across the leaflet. Black points are representative images on the left. Data collected from 2 animals per condition, 6-8 leaflets per animal. (Unpaired t-test: p = 0.4295) I. Left: Representative superficial layer images of a P8 WT and β2 KO retina flat mounts stained for endothelial cell marker CD31. Scale bar = 500μm. Right: Average vessel density across each leaflet at P8. Each point represents 1 leaflet. Points are acquired by averaging all 100μm windows across the leaflet. Black points are representative images on the left. Data collected from 2 animals per condition, 6-8 leaflets per animal. (Unpaired t-test: p = 0.6603) J. Density of vessels in center and peripheral retina at the superficial (orange) and deep (teal) layers as a function of age in WT and β2 KO mice. Central region 0.4mm from the optic nerve (solid line in E). Peripheral region 1.6mm from the optic nerve (dashed line in E). (Linear mixed effects model center superficial: p = 0.180, peripheral superficial: p = 0.645, center deep: p = 0.398, peripheral deep: p =0.399) K. Representative image of superficial layer tip cell (ESM1) and vasculature (CD31) at P5. Yellow dotted line is the angiogenic front designated as 150μm from the leading edge of growth. Inset: thresholded image with yellow arrows denoting ESM1 signal localized to the vasculature. Scale bar is 50μm. L. Quantified ESM1 signal normalized to angiogenic front vessel area in WT and β2 KO. Black point is the representative image in E. Summary statistics are median and standard deviation. Data collected from 3 animals per condition, 3-5 fields of view per mouse. (Mann-Whitney U test P3: p = 0.5313, P5: p = 0.7322)

The role of neural activity in retinal angiogenesis has been supported by multiple lines of evidence and appears to be layer-specific. Dopaminergic signaling has been shown to inhibit growth of the superficial vasculature (Liang et al. 2023), whereas cholinergic activity is required for proper development of the deep vascular layer (Weiner et al. 2019). In addition, glutamatergic signaling promotes deep layer angiogenesis by inducing the expression of pro-angiogenic factors in Müller glia (Biswas et al. 2024a). Together, these findings suggest that neural activity influences vascular development through both direct and glia-mediated mechanisms.

In the retina, there are two types of glia: astrocytes, which are restricted to the superficial retina, and Müller glia, which span the full thickness of the retina and have lateral processes that extend into the two synaptic layers of the retina. The superficial vasculature is closely associated with the endfeet of Müller glial and the astrocytes that reside on the surface of the retina (Gerhardt et al. 2003), while subsets of Müller glia lateral processes form a near-complete ensheathment of deeper capillary beds as well as the diving vessels that connect them (Grimes et al. 2024). Retinal vasculature emerges in the first two postnatal weeks, prior to visual experience. During this time, the immature retina exhibits spontaneous activity in the form of propagating retinal waves, and the Müller glia lateral processes are stabilizing in the extracellular space (Tworig et al. 2021).

Growing evidence suggests that glial cells are important for vascular development in the brain (Ma et al. 2024; Minocha et al. 2015; P. Wang et al. 2024) and in the retina (Biswas et al. 2020; Bose et al. 2025). Glial contributions to retinal angiogenesis have largely been attributed to the secretion of pro-angiogenic factors (Rattner et al. 2019; Biswas et al. 2024). However, direct physical interactions between glia and endothelial cells may also provide instructive cues. For example, superficial vasculature growth is disrupted when astrocyte patterning across the surface of the retina is perturbed (Gerhardt et al. 2003; O’Sullivan et al. 2017). In contrast, whether contact-mediated signaling contributes to the patterning of the deeper retinal vasculature remains unknown. With lateral processes in the deeper vascular layers and stalks spanning the full retina thickness, Müller glia are well positioned to influence the patterning of these deeper vascular beds. Moreover, developing Müller glia exhibit calcium transients in response to neurotransmitters released during retinal waves (Rosa et al. 2015; Tworig et al. 2021). Whether such signaling occurs at Müller glial endfeet surrounding developing vessels remains unknown.

To investigate how neuro-glial-vascular interactions contribute to retinal angiogenesis, we used the following approach. First, we characterized the coordinated development of the three vascular layers in the retina and demonstrated that this pattern is maintained in the absence of cholinergic retinal wave activity. Second, we established the developmental timeline of Müller glia-vasculature associations. Finally, we used two-photon calcium imaging to test whether the Müller glia endfeet associated with blood vessels exhibit compartmentalized calcium transients, providing initial evidence for active signaling between Müller glia and vasculature during development.

## Methods

### Animals

Mice of both sexes, ranging from postnatal day 5 to 70, were used in this study and housed under a 12-h light/dark cycle. All animal procedures were approved by the University of California, Berkeley Institutional Animal Care and Use Committee and conformed to the National Institutes of Health Guide for the Care and Use of Laboratory Animals, the Public Health Service Policy, and the Society for Neuroscience Policy on the Use of Animals in Neuroscience Research.

### Immunohistochemistry

Eyes were fixed in 4% PFA in 1x PBS for 1.5-2 hours at 4C, rinsed 3x with 1x PBS and stored at 4C until use. Stored eyes were then dissected to isolate the retina and blocked for 1-3 hours on a shaker in a 1x PBS solution with 0.3% Triton X-100 and 3% normal donkey serum (NDS). Primary antibodies were diluted in the same solution and applied for 5-7 days on a shaker at 4C. In figure 1, retinas were incubated with a rat-anti-CD31 antibody (BD Biosciences (550274), 1:100) for blood vessels and goat anti-ESM1 antibody (R&D Systems, 1:100) for tip cells. In figure 2, retinas were incubated with rabbit anti-EAAT1 (Genetex, GTX134060; 1:250) to label Müller glia. In figure 3, retinas were incubated with mouse anti–AQP4 (clone AG4.1; Santa Cruz Biotechnology; 1:200) to label glial endfeet. Secondary antibodies (CD31: Alexa-Fluor 568 and 647 donkey-anti-rat 1:500, ESM1: Alexa-Flour 568 donkey-anti-goat 1:500, EAAT1: Alexa-Fluor 647 donkey-anti-mouse 1:500, AQP4: Alexa-Fluor 488 donkey-anti-rabbit 1:500) were diluted in PBS and Triton and retinas were incubated for 24 hours.

**Figure 2:**
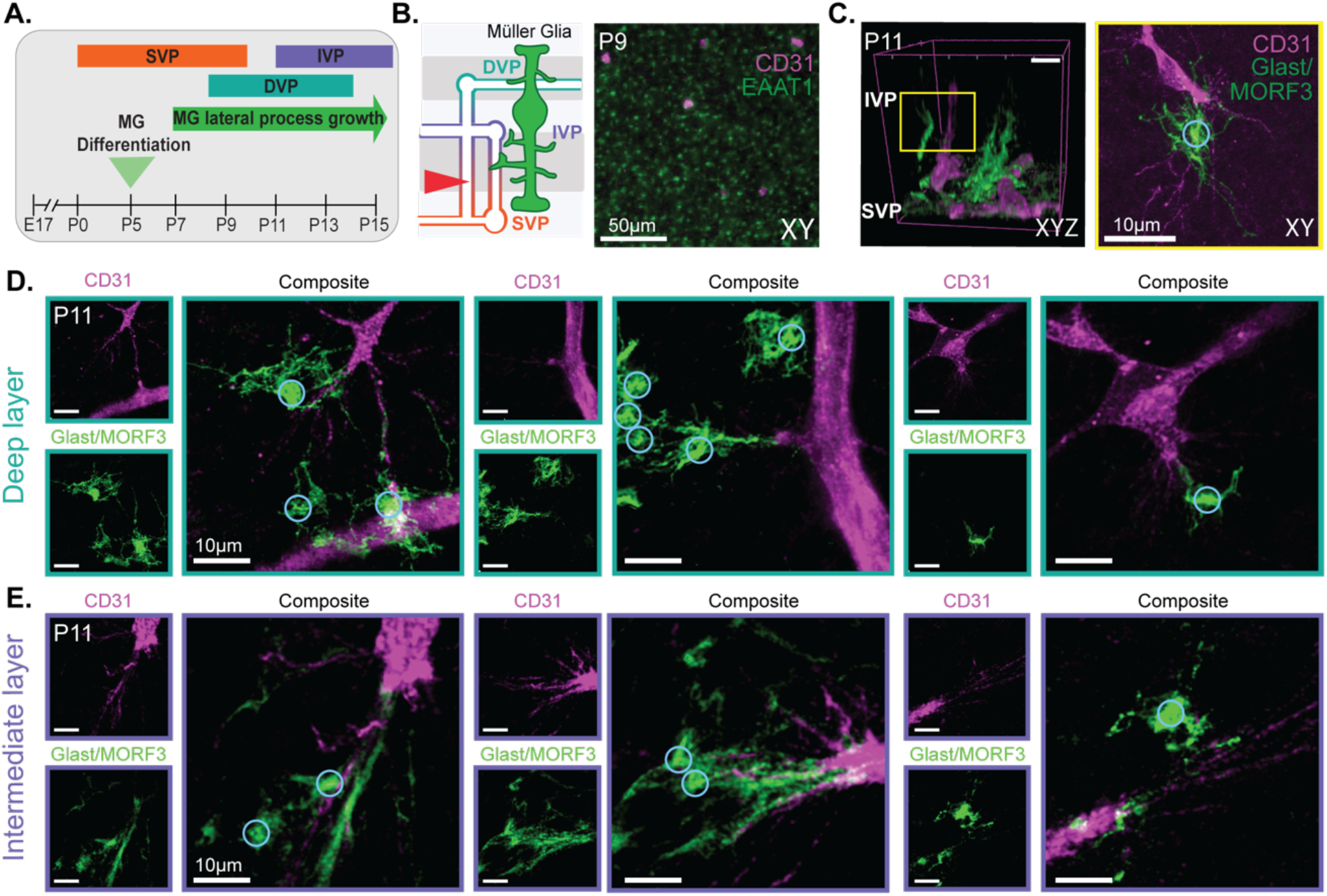
Müller Glia processes interact with active sites of growing vasculature. A. Timeline of vasculature and Müller glia development prior to eye opening. B. Left: A cross-section view of the retina: glial cells (green) and layers of vasculature (orange, purple, teal). Red arrow denotes layer of retina visualized on the right. Right: Confocal image of developing IPL. Müller glia stalks are labeled with an antibody to EAAT-1 (green). Diving vessels labeled with an antibody to CD31 (magenta). Scale bar = 50μm C. Left: 3D reconstructions of endothelial tip cell and lateral processes of Müller glia extending in the IPL at P11. Vasculature labeled with an antibody to CD31 (magenta), Müller glia are sparsely labeled using a MORF stochastic reporter (green), glia stalk circled in blue. Right: XY Confocal image of yellow box. Scale bar = 10μm. D. Müller glia lateral process and tip cell contact point examples at the deep layer at P11. Glia stalk circled in blue. Scale bar = 10μm. E. Müller glia lateral process and tip cell contact point examples at the intermediate layer at P11. Glia stalk circled in blue. Scale bar = 10μm.

**Figure 3:**
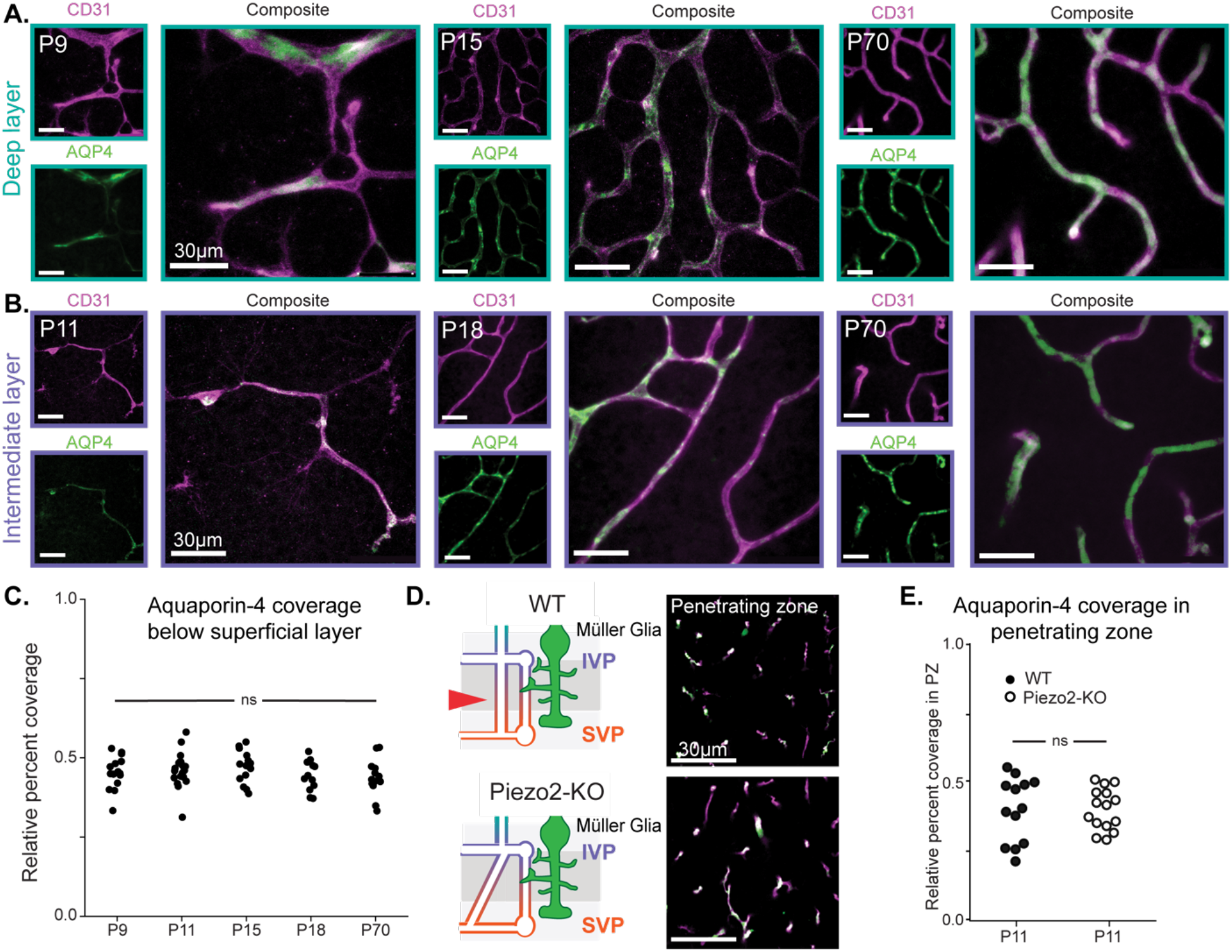
Early establishment of developed Müller glia endfeet coverage. A. Example images of deep layer vasculature throughout development and through adulthood. Vasculature labeled with an antibody to CD31 (magenta). Müller glia endfeet are labeled with an antibody to aquaporin-4 (AQP4, green). Scale bar = 30μm. B. Example images of intermediate layer vasculature throughout development and through adulthood. Vasculature labeled with an antibody to CD31 (magenta). Müller glia endfeet are labeled with an antibody to aquaporin-4 (green). Scale bar = 30μm. C. Quantification of relative percent coverage of aquaporin-4 of the vasculature from right below the superficial layer through the deep layer for A-B. Data collected from 3 animals per age, 5 images per animal. (One-way ANOVA, p = 0.36) D. Left: Schematic of aberrant vessel growth in Six3/Piezo2 KO compared to wildtype. Right: Example images of a P11 retina in wildtype and Six3/Piezo2 KO conditions. Penetrating zone (PZ) determined by capturing 10um below the superficial layer. Vasculature labeled with an antibody to CD31 (magenta). Müller glia endfeet are labeled with an antibody to aquaporin-4 (green). Scale bar = 30μm. E. Quantification of D. Relative percent coverage of aquaporin-4 on the vasculature of only the penetrating zone. Data collected from 3 animals per condition, 5 images per animal. (Unpaired T-test, p = 0.07)

### Sparse labelling of Müller glia

To sparsely label Müller glia, GLAST-Cre;MORF3 mice were generated by crossing GLAST-CreER (JAX strain 012586) mouse with Cre-ERT driven by the endogenous *Slc1a3* promoter to a MORF3 mouse (JAX strain 035403,Veldman et al. 2020) that uses mononucleotide repeat frameshift as a translation switch for cell labelling in vivo. Cre-mediated recombination was induced via intraperitoneal injection of 4-hydroxytamoxifen (50:50 E and Z isomers, Sigma-Aldrich) dissolved in sunflower seed oil. Injections of 0.5mg tamoxifen were made 2 and 4 days before each experiment.

Retinas were fixed and stained following the same immunohistochemistry protocol as above. MORF3 mice express a Cre-dependent tandem fluorescent protein with 20 V5 epitopes that switches on when the Cre has recombined the protein. To label recombined Müller glia, retinas were incubated with a chicken-anti-V5 primary antibody (Fortis Life Sciences, 1:200), and a goat-anti-chicken Alexa Fluor 488 (1:1000) secondary antibody.

### Confocal imaging of whole mount retinas

Retinas were flat mounted and imaged using confocal microscopy. To capture the entire retina leaflet, we used a 6×6 20× tile scanning imaging protocol. Leaflet images were oriented to have the optic nerve on the left and the periphery on the right. The sliding 100μm window began at the optic nerve and expanded and moved vertically to the periphery of the retina. Within each window, vessels were skeletonized to generate total vessel length within that region and normalized to the size of the window. Samples were imaged using a tile-scanning confocal microscope [Zeiss LSM 880 NLO OPO] to obtain 3D reconstructions of the tissue. The axial position of the three vascular layers will be mapped and segmented. Each vascular layer will be processed to remove background fluorescence and then manually thresholded to generate a mask.

For assessing tip cells at the superficial layer, four to five single 20x images were acquired per animal at the angiogenic front in whole-mount retinas co-labeled for vasculature (CD31) and tip cells (ESM1). For glia endfoot vascular coverage, single 20x vasculature z-stacks were taken in whole-mount retinas co-labeled for vasculature (CD31) and glia endfeet (AQP4) just below the superficial layer and through the deep layer.

For sparsely labeled glial imaging, a similar tile-scanning approach was used on a confocal microscope [Zeiss LSM 980 NLO FLIM system] equipped with a 63x oil-immersion objective. To resolve fine glial lateral processes and endothelial tip cell filopodia while avoiding saturation of glial stalks and blood vessels, high dynamic range (HDR) imaging was employed using ZEN software.

### Image analysis of AQP4 and ESM1 coverage with the vasculature

To quantify the extent of glial endfoot associations with retinal blood vessels, confocal Z-stacks of AQP4 (endfeet) and CD31 (vasculature) immunofluorescent images were processed using ImageJ/FIJI. Each channel was first filtered with a 3D median filter and then collapsed into a 2D image using a sum intensity projection. To quantify AQP4 specifically localized to the vessels, a binary mask was generated from the vasculature channel and applied to the AQP4 channel. Integrated density was then calculated from the overlap image. Relative AQP4 coverage was computed by dividing the integrated density of the overlap image by the integrated density of the vasculature projection. This ratio, expressed as a percentage, represents the relative percent coverage of glial endfeet on the retinal vascular bed.

For ESM1 vascular coverage, we used a similar approach. To isolate the angiogenic front, ROIs were manually drawn spanning 150-200μm from the leading edge of vascular growth towards the established plexus. Within each ROI, a binary mask of the vasculature was generated from the CD31 channel to restrict analysis to intravascular pixels. ESM1 fluorescence intensity was measured within the mask. ESM1 fluorescence values were normalized to the vessel area within the angiogenic front ROI, generating a measure of ESM1 signal per unit vasculature area.

### Image processing to visualize glial cell/vasculature morphology

Dual-channel confocal z-stacks were acquired at 63× magnification. Images were processed using 3D median filtering, multi-scale edge enhancement, and Noise2Void deep learning denoising (Krull et al. 2019). Noise2Void models were trained independently for each channel, then applied to denoise full volumes. Denoised stacks were max-projected in FIJI/Image J for visualization.

To generate 3D reconstructions of example glial cells, which were sparsely labeled in MORF mice, and for vasculature, which was labeled with CD31, *Z* stacks of glial cells expressing the V5 MORF protein were initially bandpass filtered in *x*–*y* space. Multiple slices at each *z*-plane were then averaged to reduce noise and enable high-quality 3D reconstruction (FIJI/ImageJ).

### Two-photon imaging

Two-photon imaging of Müller glia in the IPL was performed using a modified movable objective microscope (MOM; Sutter Instruments) equipped with an Olympus 60X, 1.0 NA, LUMPlanFLN objective (Olympus America). Two-photon excitation was evoked with an ultrafast pulsed laser (Chameleon Ultra II; Coherent) tuned to 810 nm. Laser power ranged from 7 - 12 mW for all recordings. The microscope was controlled by ScanImage software (www.scanimage.org). Scan parameters were [pixels/line x lines/frame (frame rate in Hz)]: 256 x 256 (2.96), at 1–2 ms/line. When imaging fluorescence in Cal520-labeled Müller glial stalks, processes, and endfeet, the focal plane was located within half the distance from the ganglion cell layer to the inner nuclear layer.

### Electrophysiological Recordings

Whole-cell voltage clamp recordings were made from whole-mount retinas while simultaneously imaging Cal520 fluorescence. Under infrared illumination, RGC somas were targeted for voltage clamp recordings using glass microelectrodes with a resistance of 4-7 MΩ (PC-10 pipette puller; Narishige) filled with an internal solution containing [in mM] 98.3 K-gluconate, 40 HEPES, 0.6 EGTA, 1.7 KCl, 5 MgCl2, 0.3 Na3GTP, 2 Na2ATP (pH 7.2 and 290 mOsm). Signals were acquired using pCLAMP10 recording software and a MultiClamp 700A amplifier (Molecular Devices), sampled at 10 kHz and low-pass filtered at 1 kHz. Custom MATLAB scripts generated TTL pulses to simultaneously trigger gap-free acquisition in pCLAMP and image acquisition in ScanImage, ensuring precise temporal alignment between EPSCs and fluorescence signals.

Retinal waves were identified as negative-going current deflections (troughs). To detect these events, the filtered trace was inverted and local maxima were identified using MATLAB’s findpeaks function. A user-defined minimum peak prominence ensured that detected events exceeded baseline noise fluctuations, while a user-defined inter-event distance prevented multiple detections within a single wave event. Manual inspection was done to ensure removal of noise artifacts and inclusion of visually confirmed wave-associated deflections. The charge of EPSCs was calculated by integrating current over time using the MATLAB trapz function (which uses trapezoidal numerical integration to calculate the AUC) following baseline subtraction.

### Pharmacology

For pharmacology experiments, after 5-10 minutes of recording data in ACSF, pharmacological agents were added to the perfusion, and experimental recordings were obtained 2-4 minutes afterwards. Gabazine (5 µM, Tocris) was prepared in oxygenated ACSF.

### Image processing and analysis: calcium imaging

All images were processed using custom scripts in FIJI/ImageJ (National Institutes of Health) and MATLAB R2025b (MathWorks) (Schindelin et al. 2012). For calcium imaging movies, following non-rigid motion correction, the Astrocyte Quantitative Analysis (AQuA) software identified regions in the recording field of view (FOV) that exhibited changes in fluorescence (Pnevmatikakis and Giovannucci 2017; Wang et al. 2019). Fixed size (2 µm x 2 µm) regions of interest (ROIs) were then manually placed over these regions and labeled as either stalks, lateral processes, or endfeet. Stalks were defined as semiregularly spaced, punctate regions of fluorescence 2-3 µm in diameter in average intensity projection images. Endfeet were defined as ROIs located within 2 µm of a blood vessel, typically directly overlapping the vessel. Lateral processes were defined as ROIs that did not overlap with stalk ROIs and were not within 2 µm of a blood vessel. ROIs 5 µm from the borders of the FOV were excluded from analysis. Stalk – endfoot pairs were classified as such only when a contiguous anatomical connection between compartments was visually confirmed.

Fluorescence intensity is reported as the average intensity across all pixels within the area of each ROI, and normalized as the relative change in fluorescence (ΔF/F) as follows:

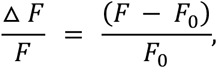

where F is the instantaneous fluorescence at any time point and F_0_is the baseline fluorescence, defined as the median fluorescence value over the duration of the trace.

ΔF/F traces were smoothed using a 3-frame median filter. Traces were Z-scored relative to the mean and standard deviation of the full recording, and calcium transients were defined as events exceeding a Z-score threshold of 2.5. Each recording was visually inspected to ensure accuracy of peak detection. Artifacts arising from baseline drift were removed and peaks not originally defined as events were included. A 2-second refractory period was imposed for event detection to prevent double-counting of individual (sustained) events. Calcium transients were classified as correlated if they occurred 1 second prior to or 5 seconds following the peak of a wave-associated EPSC (based on observed temporal relationships). Event frequency was defined as the number of detected calcium transients divided by the recording duration. The proportion of calcium transients correlated with waves was defined as the number of correlated transients divided by all transients in the recording. Calcium transients in stalk-endfoot pairs were deemed correlated if they occurred within ± 3 seconds of a calcium transient in their paired compartment.

To test whether calcium events were temporally correlated with retinal waves beyond chance levels, we used a permutation test for each ROI. First, we computed the observed fraction of calcium events classified as correlated. We then generated a null distribution by block-shuffling event timestamps within each recording (1,000 permutations) while preserving the number and temporal structure of events and recomputed the fraction of correlated events for each permutation. An ROI was classified as significantly correlated if its observed fraction exceeded the 95th percentile of the null distribution.

### Statistical Analyses

All statistical analyses were performed in Python. For two-sample comparisons, data were first assessed for normality using the Shapiro-Wilk test. For independent two-sample comparisons, normally distributed data were tested for homogeneity of variance using Bartlett’s test and analyzed using a two-tailed Student’s *t*-test. When data were not normally distributed, differences were assessed using the nonparametric Mann-Whitney U test (Wilcoxon rank-sum test). Comparisons among three or more groups were evaluated using one-way ANOVA. Vessel density in the center and periphery of the retina in WT and β2 KO mice across age was analyzed using a linear mixed-effects model. Significance was defined as p <0.05. Summary statistics are reported as mean ± SEM for all unless otherwise noted.

## Results

### Retina vasculature grows uniformly and independent of cholinergic retinal waves

We first assessed the impact of disrupting cholinergic retinal waves on vasculature development. Pharmacological disruption of cholinergic activity significantly affected the maturation of the deep vasculature (Weiner et al. 2019), whereas a genetic manipulation had a smaller impact (Biswas et al. 2024a).

To characterize the development of the retina vasculature, we performed immunohistochemistry for the endothelial cell marker CD31 (Toma et al. 2024) in flat-mounted retinas isolated from wild-type mice from postnatal day 5 (P5) to eye opening at postnatal day 14 (P14; Fig.1B,C). Samples were imaged using tile-scanning confocal microscopy to obtain 3D reconstructions, and the axial positions of the superficial, intermediate and deep vascular layers were segmented. To describe the maturation of the vasculature, we quantified vessel density as the total vessel length per unit retinal area using a sliding 100-micron window from the optic nerve head to the distal end of a retina leaflet, allowing direct comparison of central and peripheral regions.

Consistent with the previous description of retinal angiogenesis (Krimpenfort et al. 2023; Milde et al. 2013; Darche et al. 2022), the superficial vasculature emerged first and expanded centrifugally from the optic nerve, followed by a period of pruning in the second postnatal week (Fig. 1D-G). Vessels from the superficial and deep layers grow perpendicularly into the tissue and meet to establish the intermediate layer. While the superficial and diving vessels exhibited radial outgrowth, the intermediate layer developed a more discontinuous, patch-like pattern (Fig. 1E).

To study the impact of cholinergic activity on vasculature maturation, we isolated retinas from mice lacking the β2 subunit of nicotinic acetylcholine receptors (β2-nAChR knockout mice), which have dramatically reduced cholinergic wave activity during the first postnatal week (Bansal et al. 2000). At both P5, when the superficial vasculature has extended approximately halfway across the retina, and P8, when wave activity recovers in β2–nAChR knockout mice, we found that vascular density remained comparable to wildtype (Fig. 1H–I), consistent with prior reports (Biswas et al. 2024). We found that in both superficial and deep layers vascular growth progressed normally despite the reduction in cholinergic wave activity (Fig. 1J).

Although overall vascular density was unchanged in β2–nAChR knockout mice, previous studies have shown that dopaminergic activity can influence angiogenesis by regulating tip cell sprouting during the same developmental time window when cholinergic retinal waves occur (Liang et al. 2023). Tip cells are specialized endothelial cells that extend filopodia at the leading edge of vessels to guide angiogenesis. To determine whether reduced cholinergic waves affected tip cell sprouting, we labeled superficial layer tip cells with an antibody against tip cell marker ESM-1 and quantified ESM-1 signal during superficial layer angiogenesis (Fig. 1K). ESM-1 signal was normalized to the vessel area comprising the angiogenic front, defined as approximately 150μm from the leading edge of vascular growth. At both P3 and P5, ESM-1 expression was unchanged in β2–nAChR knockout mice (Fig. 1L). Taken together, these findings suggest that signaling from neurons modulating tip cell sprouting and formation of the superficial, deep, and intermediate vascular layers proceed normally despite reductions in cholinergic wave activity.

### Müller glia associate with actively growing vasculature and establish early endfoot coverage

Next, we established the stage of development when Müller glia – vasculature contacts first appear. Subsets of Müller glia lateral processes form specialized contacts with the vasculature. These contacts occur at endfeet which are specialized compartments of the glial lateral process that interface with blood vessels. During angiogenesis in the deeper vascular layers, Müller glia lateral processes emerge and are highly motile (Fig. 2A) (Tworig and Feller 2022; Tworig et al. 2021).

To determine when the Müller glia–vasculature interface first emerges, we immunolabeled the vasculature (CD31) and Müller glia (GLAST) during angiogenesis of the deeper layers. Before Müller glia fully elaborate their lateral processes, diving vessels below the superficial layer grow near numerous Müller glial stalks (Fig. 2B). With this labeling method, we were unable to resolve individual Müller glia within this densely populated region. To gain single-cell resolution, we used a Cre-dependent stochastic mononucleotide frameshift mutation mouse (GlastCre:MORF3). This method sparsely labels Müller glia, allowing us to visualize individual glial cells contacting blood vessels. To identify sites of active vascular growth, we focused on endothelial tip cells. We observed close associations between Müller glial lateral processes and endothelial tip cells in both the deep and intermediate vasculature at P11, when both layers are actively growing (Fig. 2C–E).

To determine whether these early vascular associations reflect mature glial endfeet, we immunolabeled the vasculature (CD31) and Aquaporin-4 (AQP4). AQP4 is a water channel protein enriched at Müller glial endfeet that contact retinal blood vessels, where it contributes to water homeostasis, and is widely used as a marker for vascular-contacting glial endfeet (Nagelhus et al. 1998) (Fig. 3A-B). In the retina, AQP4 immunoreactivity was restricted to Müller glial processes at vascular contact sites, with little to no detectable labeling along nonvascular regions of the glial membrane. Notably, AQP4 labeling was spatially heterogeneous along the endfoot, consistent with prior observations that AQP4 is enriched in subdomains rather than uniformly distributed across the entire endfoot membrane. At every layer and developmental stage examined, the extent of AQP4 coverage was comparable to that observed in adults (Fig. 3C).

We examined the robustness of these associations between glia and retina vasculature and repeated these experiments in a mouse model in which vasculature growth is perturbed. Piezo2 in retinal neurons regulates diving vessel trajectory; loss of Piezo2 disrupts normal patterning of diving vessels, leading superficial vessels to invade the IPL along aberrant, diagonal paths instead of their typical perpendicular descent (Toma et al. 2024)(Fig. 3D). We measured the relative percent AQP4 coverage within the penetrating zone (PZ)—the region spanning the diving vessels between the superficial and deep layers. AQP4 coverage in the PZ was unchanged in mice lacking Piezo2 (Fig. 3E). These findings indicate that Müller glia dynamically follow and establish endfoot contacts with growing vessels during the earliest stages of angiogenesis even when vessel growth trajectories are disrupted.

### Calcium signaling in Müller glial endfeet is compartmentalized and occur independently of retinal waves

Having established that Müller glial endfeet form early vascular contacts, we next examined whether these specialized compartments exhibit calcium transients that are temporally correlated with retinal waves. To assess this, we conducted two-photon calcium imaging of Müller glial stalks, processes, and endfeet while simultaneously performing voltage-clamp recordings from retinal ganglion cells (RGCs). Retinas from wildtype mice (age P9-12) were bath-loaded with the calcium indicator Cal520-AM, which preferentially labels Müller glia (Rosa et al. 2015; Uckermann et al. 2004). Imaging fields were centered on diving blood vessels, typically within the ON sublamina of the inner plexiform layer. Regions of interest were defined by pixels exhibiting correlated increases in fluorescence during the recording window.

We could reliably distinguish Müller glial stalks, lateral processes not associated with blood vessels, and endfeet, defined as lateral processes closely opposed to vessels (Fig. 4A, B). Along individual blood vessels, multiple endfeet exhibited calcium transients that were spatially restricted and temporally independent, and in some cases, could be traced to a nearby stalk (Fig. 4B). All three Müller glial compartments – stalks, lateral processes and endfeet – exhibited spontaneous calcium transients at comparable frequencies (∼0.5 – 3 events per minute; Fig. 4D, E). In each compartment, a subset of events was temporally correlated with retinal waves, defined as ΔF/F transients occurring within 5 seconds of a wave (Fig. 4F), as well as spontaneous events not time-locked waves, consistent with previous reports (Rosa et al. 2015; Tworig et al. 2021).

**Figure 4.**
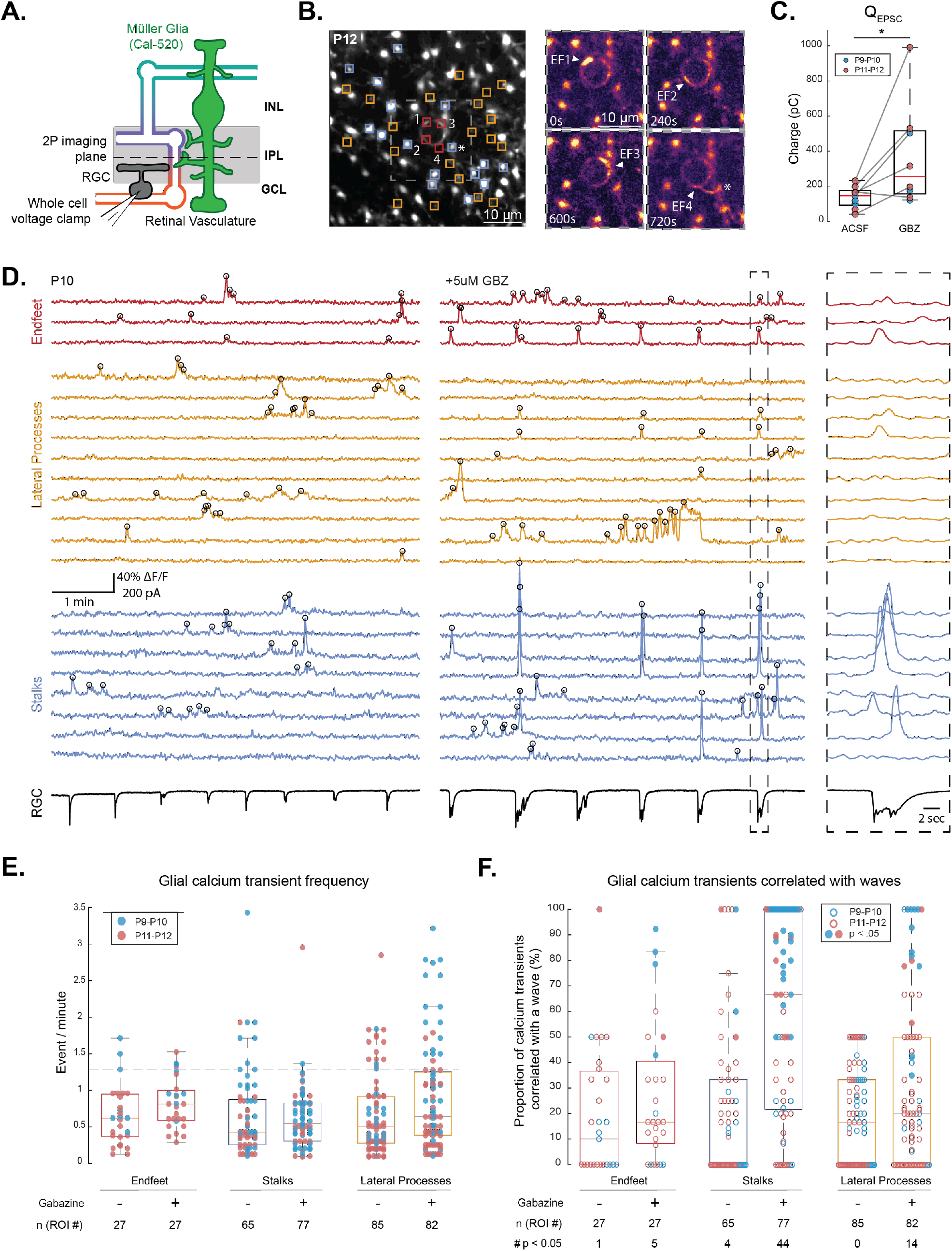
Müller glia display compartmentalized spontaneous and wave-correlated calcium transients during development. A. Diagram of experimental setup; Müller glia in whole-mount retina were bath-loaded with Calbryte-520 AM (green) and imaged with a two-photon microscope at the depth of the inner plexiform layer (IPL; dashed line). Fields containing a diving blood vessel were selected for recording. Retinal waves were detected via simultaneous voltage-clamp recordings of excitatory postsynaptic currents (EPSCs) from a retinal ganglion cell (RGC) held at –60 mV. B. Left, Average projection of a field of view (FOV) showing Cal520-labeled Müller glia and a diving blood vessel. Regions of interest (ROIs) are overlayed for stalks (blue), lateral processes (yellow), and endfeet (red) that exhibited calcium transients during recording. Asterisk indicates a stalk with an observable endfoot. Right, average intensity projections of the boxed region over separate 30-second periods surrounding events (start time indicated). Scale bars = 10 µm. C. Summary plot of total integrated EPSCs (Q_epsc_) in control (ACSF) and gabazine (GBZ, 5 µM). Each data point represents mean Q_epsc_ per recording (n = 101 from 9 cells and 139 EPSCs from 8 cells for control and gabazine conditions). *p < 0.05, two-tailed Student’s *t*-test. D. Example ΔF/F traces from Cal520-labeled Müller glia endfeet (red), lateral processes (yellow), stalks (blue), and wave-associated EPSCs recorded from an RGC held at -60 mV (black). Traces are before and after gabazine (5 µM) application. Right, Müller glial calcium transients during a retinal wave, expanded in time to highlight differences across compartments. E. Summary plot of calcium event frequency across Müller glia compartments in control and gabazine. Each data point represents an ROI. Dashed line indicates mean EPSC frequency across conditions (1.35 events/minute). F. Summary plot of correlations of glial calcium transients and retinal waves in control and gabazine. Each data point represents an ROI. Shaded points are considered significantly correlated according to a permutation test: numbers at the bottom list the number of data points whose correlations ranked higher than 95% of the permuted correlations for that ROI. Boxes represent interquartile range (IQR) with median (red line); whiskers extend to 1.5 X IQR. Data collected from 6 animals over 11 recordings.

To test the effects of neurotransmitter spillover during waves on Müller glial calcium signaling, we bath-applied the GABA-A receptor antagonist gabazine (5 µM), which increased the integrated current associated with wave-induced compound EPSCs, consistent with elevated excitatory neurotransmitter release (Fig. 4C). Gabazine led to a significant increase in the proportion of stalk calcium transients that correlated with retinal waves and the proportion of stalks exhibiting correlative calcium transients within a given field of view (Fig. 4F, S1). However, gabazine did not have a significant effect on the correlation of calcium transients in endfeet with waves (Fig. 4F). These findings indicate that neurotransmitter spillover during retinal waves is not the primary driver of calcium transients in Müller glial endfeet.

We next examined whether calcium transients exhibit compartment-specific dynamics. First, we observed that wave-triggered averages of wave-correlated calcium transients had distinct temporal profiles in stalks compared with lateral processes and endfeet (Fig. S2A). We next examined calcium transients within individual stalk-endfoot pairs. Pairs were identified as active endfoot regions contiguous with a nearby stalk (Fig. S3A), structures that were comparable to stalk-endfoot pairs observed by immunolabeling (Fig. S3B). Comparison of ΔF/F traces revealed substantial heterogeneity in wave correlations and cross-compartment correlations across pairs (Fig. S3C–E). In all pairs, there were calcium transients that occurred independently of the paired compartment for either the stalk, the endfoot, or both. In 7/12 pairs, there was no correlated calcium activity across compartments, and in 3/12 pairs, there were no detectable calcium transients in the stalk from which the endfoot derived.

Together, these data indicate that Müller glial endfeet exhibit compartmentalized calcium transients during development. Although a subset of endfoot transients were temporally correlated with retinal waves, enhancing neurotransmitter spillover had minimal impact on the prevalence of this correlation, suggesting that calcium signaling at the glial-vascular interface is highly independent of spontaneous neuronal activity.

## Discussion

Here, we have described a close association between Müller glia and the vasculature throughout retinal angiogenesis. Lateral processes from Müller glia formed contact points with endothelial tip cells – the leading edge of growing vasculature. Müller glia also established AQP4-enriched endfoot contacts with growing vessels from the earliest stages of angiogenesis, including in mice with aberrant vascular trajectories. Finally, Müller glial endfeet exhibited robust, compartmentalized calcium signaling that was uncorrelated with the neighboring Müller glial stalks or retinal waves. Together, these findings suggest that glial-vascular interactions are not downstream of spontaneous neuronal activity but rather reflect a parallel, independent developmental program.

### Neural signaling and angiogenesis

In this study, we find that temporal progression of angiogenesis remains unchanged in retinas lacking cholinergic retinal waves. Before the onset of vision, waves are a large source of cholinergic activity in the retina. Cholinergic waves are driven by release of acetylcholine from starburst amacrine cells (SACs), and waves propagate through β2-subunit-containing acetylcholine receptors on SACs and RGCs (Bansal et al. 2000; Ford et al. 2012). However, across all three layers of vasculature, we see no change in growth when cholinergic waves are inhibited. Our findings align with previous work using the β2-nAChR knockout, which similarly disrupts cholinergic wave activity without altering angiogenesis. Interestingly in Vglut1 KO retinas, cholinergic waves persist leading to a delay in the formation of the deep vasculature (Biswas et al. 2024a). In contrast, an earlier study reported that killing starburst amacrine cells with an anti-ChAT-SAP toxin during retinal angiogenesis impaired development of the deep vasculature layer, while leaving the superficial and intermediate layers unaffected (Weiner et al. 2019). Together, these findings suggest that previously reported effects of cholinergic signaling on deep-layer angiogenesis likely occur through a mechanism independent of β2-containing nicotinic receptors and, more broadly, that cholinergic waves are not required for the overall timing of vascular layer formation.

More generally, neuronal activity has been implicated in the regulation of angiogenesis through multiple, and sometimes opposing, pathways. The primary model of angiogenesis is driven by release of vascular endothelial growth factor (VEGF) from hypoxic cells, and this process is further shaped by a variety of neuron and glia derived signals (Wälchli et al. 2023; Rattner et al. 2022). In the retina, development of the deep and intermediate vasculature depends on several pro-angiogenic factors released from neurons and Müller glia (Rattner et al. 2022, 2019). Several studies have demonstrated that angiogenic signaling pathways in the retina are influenced by neural activity (Biswas et al. 2020; Liang et al. 2023; Rao et al. 2013). For instance, cholinergic retinal waves have been reported to promote VEGF release (Weiner et al. 2019), while glutamatergic waves drive Müller glial secretion of Norrin to support deep vasculature growth (Y. Wang et al. 2012; Biswas et al. 2024a). In contrast, dopamine released from retinal neurons suppresses superficial vasculature outgrowth (Liang et al. 2023), highlighting that neural activity can both promote and restrict angiogenesis depending on the signaling context.

Whether neural activity influences angiogenesis directly through neuronal release of growth factors or indirectly via intermediary glial populations remains unresolved. This complexity is not unique to the retina but is evident across the developing CNS. In the cortex, sensory deprivation paradigms such as whisker plucking or barrel cortex lesions have been reported to alter vascular density and branching (Lacoste et al. 2014), although parallel studies reached conflicting conclusions (Whiteus et al. 2014). Together, these findings indicate that neural activity can influence angiogenesis through multiple, context-dependent mechanisms. In this context, our findings demonstrate that eliminating cholinergic retinal waves does not alter the temporal progression of angiogenesis, narrowing the conditions under which spontaneous activity is necessary for vascular development.

### Glial influences on growing vasculature

In this study, we demonstrate that Müller glia lateral processes closely associate with developing endothelial tip cells during intermediate and deep layer angiogenesis, and that AQP4 coverage remains stable at levels comparable to adulthood throughout this developmental period. Because AQP4 is selectively enriched at Müller glial endfeet that contact the vasculature, the early establishment of adult-like AQP4 coverage suggests that Müller glia develop vascular associations in tandem with, rather than after, angiogenesis. Notably, this coverage is maintained even when proper vessel patterning is perturbed, indicating that Müller glial-vasculature association is tightly coupled to angiogenesis and resilient to changes in vascular patterning. This is in contrast to findings in zebrafish, where glial coverage of the vasculature increases with development (Gall et al. 2025).

It is well established that glial cells play a functional role in vascularization. Deletion of hypoxia-inducible factor 2α from Müller glia reduces deep layer vessel density, consistent with a role for glia-derived hypoxic signaling in regulating angiogenesis (Rattner et al. 2019). Similarly, in zebrafish, glial ablation disrupts CNS angiogenesis through reduced VEGF signaling, and perturbations in gliogenesis lead to defects in vessel patterning (Matsuoka et al. 2016). Together, these findings indicate that glial populations are active participants in angiogenesis of intermediate and deep retinal angiogenesis via secretion of proangiogenic factors.

Are Müller glia instructive for growth of deep vasculature? It is well established that superficial angiogenesis is guided by an astrocytic network (Gerhardt et al. 2003), with the organization of the superficial vasculature dictated by the patterns of astrocytes (O’Sullivan et al. 2017). While we did not observe diving vessels grow along the Müller glial stalked, we did observe that Müller glial lateral processes closely associated with endothelial tip cells during angiogenesis, indicating a potential source of guidance. Ablation of Müller glia would likely compromise retinal structure given their essential mechanical and metabolic support roles (Byrne et al. 2013). Thus, approaches that could selectively disrupt Müller glial lateral process outgrowth or motility while preserving glial stalk integrity could provide a means to assess their contribution to deeper layer angiogenesis.

### Glial compartmental signaling in development and adulthood

Glial endfoot signaling at vascular interfaces is known to play crucial roles in homeostasis and neurovascular coupling in the brain (Araque et al. 2014; Díaz-Castro et al. 2023) and retina (Newman 2015). We found that Müller glial form endfeet throughout angiogenesis and that they exhibited spontaneous calcium transients. This is a period of maturation for Müller glia, as their lateral processes extend and retract across different IPL sublaminae (Wang et al. 2017; Tworig et al. 2021). Despite this dynamic remodeling, the endfeet show compartmentalized calcium transients as early as P9. The endfeet transients occur largely independent of retinal waves, even with enhanced neurotransmitter spillover via the application of a GABA-A receptor antagonist, gabazine. This contrasts with stalk calcium transients, which are more correlated with retinal waves, as previously described (Rosa et al. 2015; Zhang et al. 2019; Tworig et al. 2021). These findings are consistent with previous reports in the brain that show that endfoot calcium transients occur independent of nearby neural activity (Bonder and McCarthy 2014; Del Franco et al. 2022; Shigetomi et al. 2013).

What evokes compartmentalized calcium transients in glial endfeet? Spatially restricted endfoot calcium transients have been observed in the brain following neuromodulator application, sensory stimulation, and electrical stimulation (Mulligan et al. 2004; Montalant et al. 2024; Institoris et al. 2022; Mishra et al. 2016; Bindocci et al. 2017; Takano et al. 2006; Nizar et al. 2013). In the adult retina, endfoot calcium signals have been observed following light stimulation and ATP application (Biesecker et al. 2016; Grimes et al. 2024), though the degree of compartmentalization has not been characterized. We speculate that endfoot compartmentalization is due to differing receptor distribution and/or intracellular calcium store organization compared to other regions of the cell.

In summary, our findings show that retinal angiogenesis proceeds normally despite a dramatic reduction in cholinergic retinal wave signaling, while Müller glia establish stable AQP4-enriched endfeet on growing vessels from the earliest stages. Moreover, calcium signaling at the glial–vascular interface is largely independent of spontaneous neuronal activity, supporting a model in which Müller glia engage developing vessels through an activity-independent developmental program.

## Data Availability

The data that support the findings of this study are openly available in a repository at https://github.com/FellerLabCodeShare/Glia_Vaculature.git

The authors declare no conflicts of interest regarding this manuscript

## Acknowledgements

We thank members of the Feller lab for commenting on the manuscript. Confocal imaging experiments were conducted at the CRL Molecular Imaging Center. We would like to thank Holly Aaron and Feather Ives for their microscopy training and assistance and Etai Sapoznik for assistance in image processing. We thank Aileen Conley for assistance with generating three-dimensional reconstructions. SM supported by the University of California Dissertation-Year Fellowship. AJG is supported by the National Science Foundation Graduate Research Fellowship (DGE 2146752) and the HHMI Gilliam Fellowship (GT17844). All other authors are supported by NIH grants R01EY019498, R01EY013528, R21EY037926, and P30EY003176.

---

**Supplemental Figure 1.**
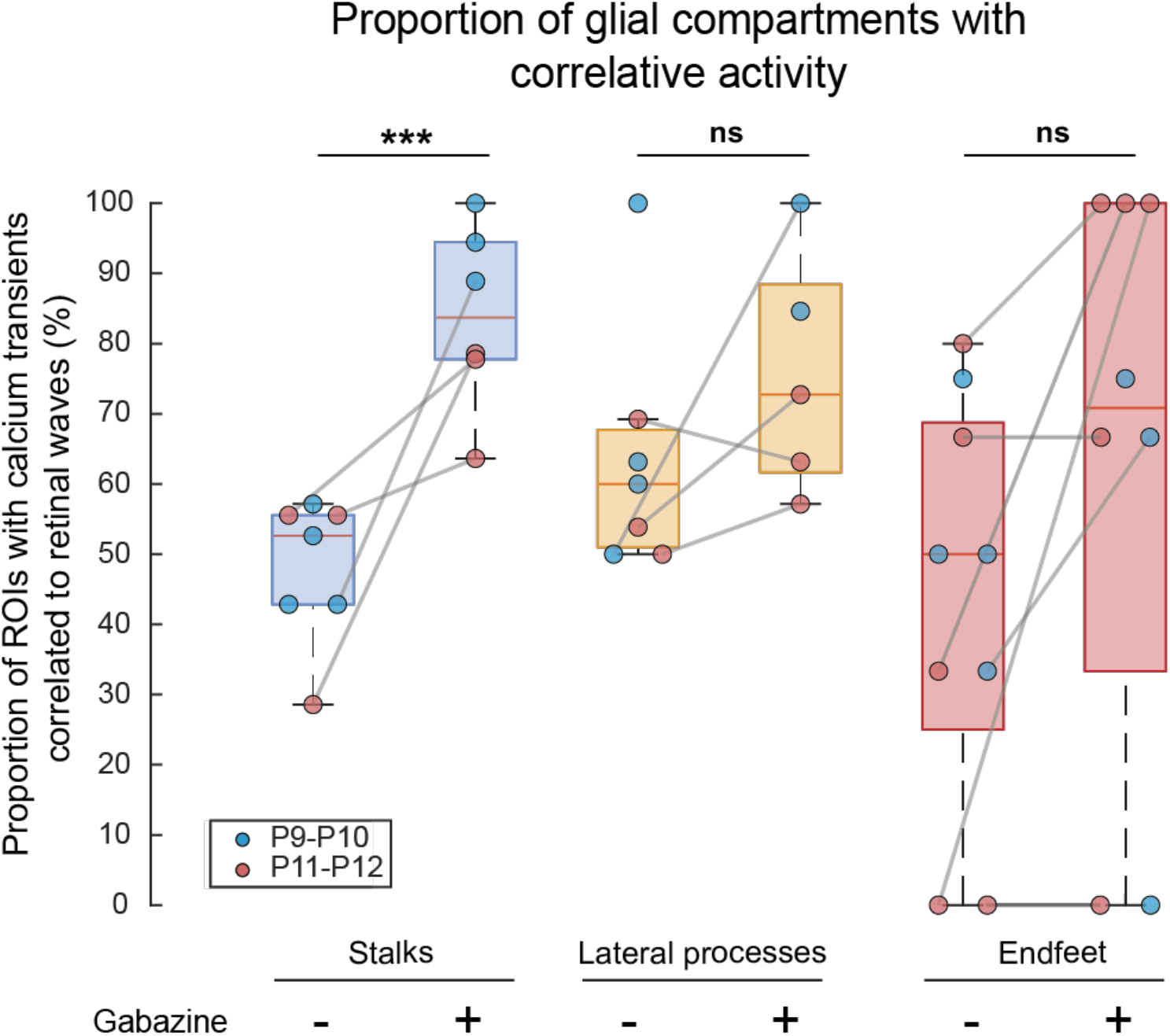
Proportion of ROIs with at least one calcium transient correlated with a retinal wave in control and gabazine (5 µM). Each data point represents the proportion of ROIs with correlative activity in a field of view. Lines indicate paired recordings before and after gabazine. ***p >.001, two-tailed Student’s t-test. Data collected from 6 animals over 11 recordings.

**Supplemental Figure 2.**
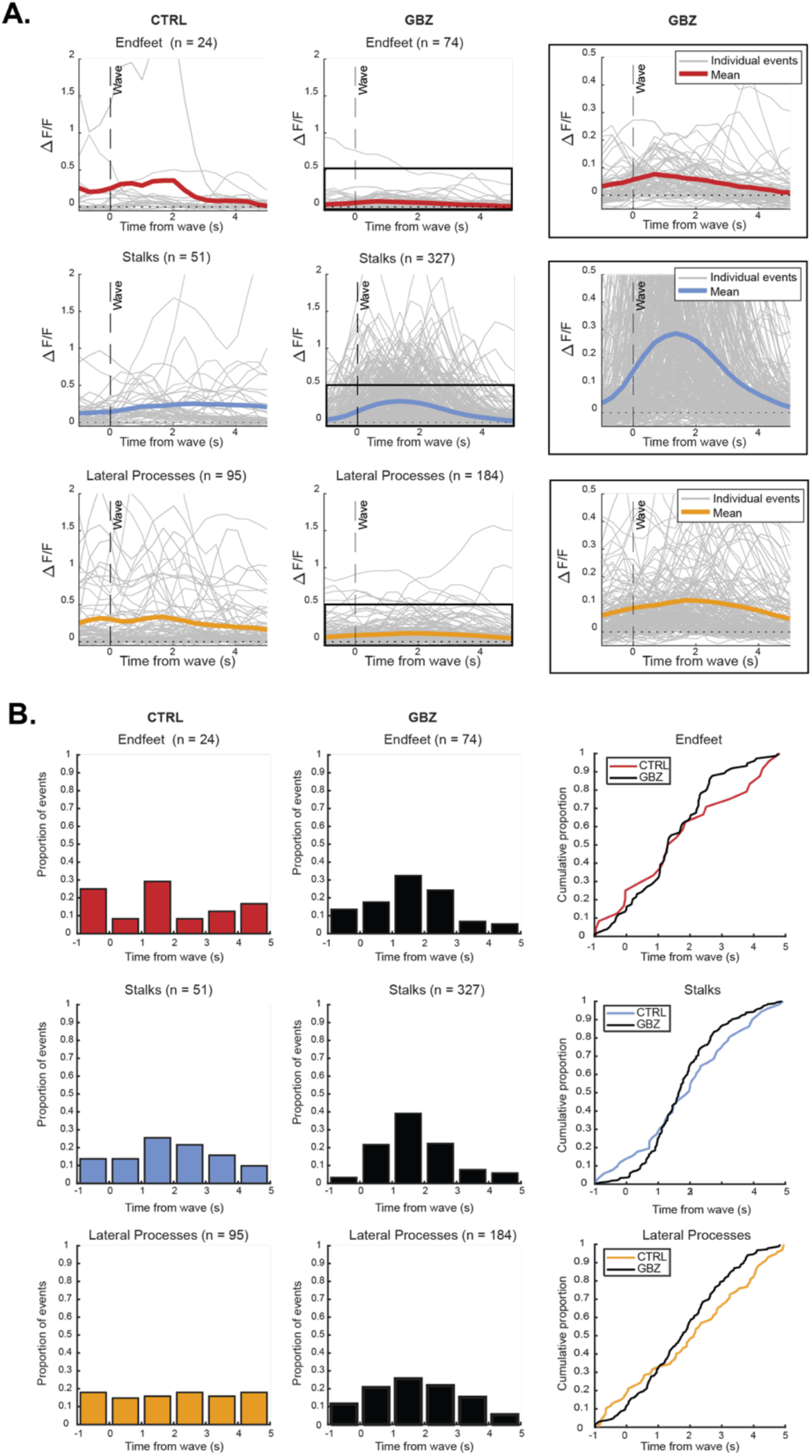
A. Left and middle, individual wave-associated calcium transients (gray) from each compartment with corresponding wave-triggered average traces overlaid (lateral processes, yellow; stalks, blue; endfeet, red) in ACSF (CTRL) and gabazine (GBZ; 5 µM). Event counts (n) are shown above each panel. Right, expanded view of the boxed region. B. Left and middle, histograms of calcium event timing relative to retinal wave peak (0 s) in ACSF and gabazine. Right, cumulative distributions of event timing across conditions.

**Supplemental Figure 3.**
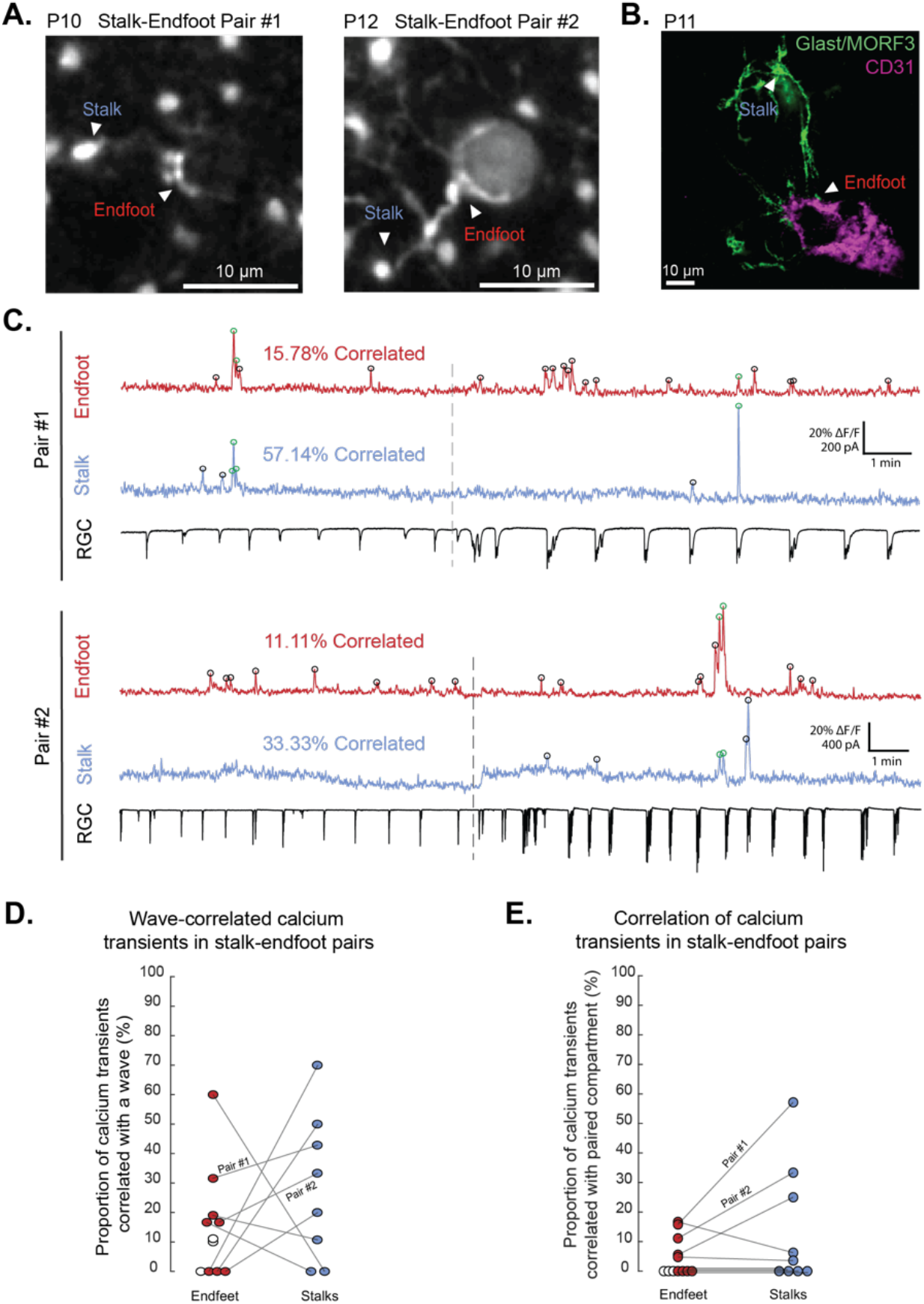
A. Representative examples of Müller glia stalks and their associated endfeet. Intensity projections from the IPL of whole-mount retina showing Cal520-labeled stalk-endfoot pairs. B. Example confocal image of growing diving vessel and Müller glia lateral processes from two individual stalks. Vasculature labeled with an antibody to CD31 (magenta), Müller glia are sparsely labeled using a MORF stochastic reporter (green). C. Example ΔF/F traces from paired endfeet (red) and stalks (blue) in A, with simultaneous voltage-clamp recordings of wave-associated EPSCs from an RGC held at -60 mV (black). Calcium events correlated between paired compartments are indicated by green circles; uncorrelated events are shown in black. Dashed lines represent gabazine (5 µM) application. D. Correlations of glial calcium transients and retinal waves in paired endfeet and stalks. Each data point represents an ROI. E. Percentage of calcium transients correlated between paired endfeet and stalks. Each data point represents an ROI. Each line connects compartments from the same cell. Open points indicate compartments with no detectable calcium transients in paired compartment. Scale bars, 10 µm. n = 127 endfoot events and 93 stalk events. Data collected from 6 animals over 8 recordings.

